# Characterization of 100 extended major histocompatibility complex haplotypes in Indonesian cynomolgus macaques

**DOI:** 10.1101/2019.12.16.878421

**Authors:** Cecilia G. Shortreed, Roger W. Wiseman, Julie A. Karl, Hailey E. Bussan, David A. Baker, Trent M. Prall, Amelia K. Haj, Gage K. Moreno, Maria Cecilia T. Penedo, David H. O’Connor

## Abstract

Many medical advancements – including improvements to anti-rejection therapies in transplantation and vaccine development – rely on pre-clinical studies conducted in cynomolgus macaques (*Macaca fascicularis*). Major histocompatibility complex (MHC) class I and class II genes of cynomolgus macaques are orthologous to human leukocyte antigen complex (HLA) class I and class II genes, respectively. Both encode cell-surface proteins involved in cell recognition and rejection of non-host tissues. MHC class I and class II genes are highly polymorphic, so comprehensive genotyping requires the development of complete databases of allelic variants. Our group used PacBio circular consensus sequencing of full-length cDNA amplicons to characterize MHC class I and class II transcript sequences for a cohort of 295 Indonesian cynomolgus macaques (ICM) in a large, pedigreed breeding colony. These studies allowed us to expand the existing database of *Macaca fascicularis* (*Mafa*) alleles by identifying an additional 141 MHC class I and 61 class II transcript sequences. In addition, we defined co-segregating combinations of allelic variants as regional haplotypes for 70 Mafa-A, 78 Mafa-B and 45 Mafa-DRB gene clusters. Finally, we defined class I and class II transcripts that are associated with 100 extended MHC haplotypes in this breeding colony by combining our genotyping analyses with short tandem repeat (STR) patterns across the MHC region. Our sequencing analyses and haplotype definitions improve the utility of these ICM for transplantation studies as well as infectious disease and vaccine research.

## Introduction

The highly polymorphic genes in the major histocompatibility complex (MHC) region are most notably involved in antigen presentation and recognition. In humans, the classical class I region contains three genes: HLA-A, HLA-B, and HLA-C. In macaques, there are only MHC-A and MHC-B orthologues, though there have been complex duplication events within this region, leading to MHC-A and MHC-B haplotypes with copy number variations that encode various combinations of major and minor transcripts as well as multiple pseudogenes (Daza-Vamenta et al., 2004; Fukami-Kobayashi et al., 2005; Watanabe et al., 2007). The class II region includes six distinct loci, which are the same in humans and macaques: DRA, DRB, DQA1, DQB1, DPA1, and DPB1 (Shiina et al., 2017). Because macaques are commonly used as models for infectious disease, vaccine, and transplant studies, an emphasis has been placed on characterizing their MHC polymorphisms (Wiseman et al., 2013; Shiina and Blancher, 2019).

Researchers using nonhuman primate (NHP) transplantation models need to control for differences in the MHC genotypes of donor and recipient tissues and often will vary degrees of disparity when testing tolerance therapies (Burwitz et al., 2017; Kean et al., 2012; Shiina and Blancher, 2019). The National Institutes of Health (NIH) has recognized the need for and importance of transplant research conducted in NHP models since many immunomodulatory strategies that promote tolerance in rodent models have failed to translate successfully to humans (Knechtle et al., 2019). This led to the creation of the Nonhuman Primate Transplantation Tolerance Cooperative Study Group, whose goals include the development of novel regimens for immune tolerance induction, as well as understanding the mechanisms responsible for induction, maintenance, and/or loss of tolerance in NHP models of kidney, pancreatic islets, heart and lung transplantation. In support of these studies, the NIH has established several breeding colonies in order to provide Indian-origin rhesus macaques, Mauritian-origin cynomolgus macaques (MCM) and Indonesian-origin cynomolgus macaques (ICM) to NHP transplant researchers. The MCM population displays extremely restricted MHC diversity with only seven ancestral MHC haplotypes (Wiseman et al., 2013). This population appears to have originated from a very small number of founding individuals who were introduced to Mauritius in the early 1500’s (Lawler et al., 1995; Tosi and Coke, 2007). While this restricted genetic diversity of the MCM population makes them extremely valuable for many types of biomedical research, it can be a disadvantage for transplant researchers whose objective is to maximize the amount of MHC disparity between donor and recipient tissues. In contrast, the ICM population displays extensive MHC diversity (Otting et al., 2012) which makes them an ideal NHP model for studies designed to evaluate the preclinical safety and efficacy of immune tolerance induction regimens that may be translated to clinical transplantation.

A number of investigators have used ICM (Ezzelarab et al., 2016; Zhang et al., 2015) as well as cynomolgus macaques from various other geographic origins (Kato et al., 2017; Matsunami et al., 2019; Shiina and Blancher, 2019) for transplantation studies. For instance, ICM were used in a study to investigate T cell and alloantibody responses with purposely mismatched donor and recipient pairs (Ezzelarab et al., 2016). Another study (Morizane et al., 2017) utilizing a Filipino cynomolgus macaque model showed that when donor and recipient are matched for MHC genotypes, there are improved outcomes with neural cell grafts. Additionally, they observed inflammation in mismatched donor-recipient pairs. This indicates that the ability to accurately and efficiently characterize MHC genetics of model species like ICM is and will remain a necessity (Shiina and Blancher, 2019).

Prior to this study, several reports have characterized ICM MHC sequence variants. In 2008, 48 novel MHC class I sequences were documented in 42 animals by Sanger sequencing of cDNA clones (Pendley et al., 2008). Likewise, Kita *et al*. (2009) surveyed 27 cynomolgus macaques of Indonesian descent and identified 34 novel *Mafa-A* sequences. Another study in 2011 (Creager et al., 2011) performed Sanger sequencing of MHC class II cDNA clones from a dozen unrelated ICM and identified 58 new transcript sequences. In the most comprehensive MHC genotyping analysis of ICM to date (Otting et al., 2012), Otting and colleagues characterized the class I and class II alleles that are associated with 32 extended MHC haplotypes in 120 ICM from a pedigreed breeding colony at the Biomedical Primate Research Centre (BPRC) in the Netherlands. More recently, these investigators evaluated MHC class II sequences in 79 ICM housed at the same colony that we examined in the present study (Otting et al., 2017). They identified 22 novel and extended 10 partial *Mafa-DQ* and *Mafa-DP* sequences. Despite these previous studies characterizing ICM MHC sequence variants, we hypothesized that many additional MHC sequences remained to be characterized in this NIH-sponsored ICM breeding colony.

Several years ago, our group introduced the use of Pacific Biosciences (PacBio) single-molecule real-time sequencing to determine full-length MHC class I transcript sequences (Westbrook et al., 2015). Rapid improvements in PacBio circular consensus sequencing technology have made it possible to generate highly accurate sequences of cDNA amplicons from MHC class I and class II transcripts that span complete open reading frames of approximately 1100bp and 800bp, respectively (Karl et al., 2017, 2014). This sequencing approach has proven useful in investigations of several understudied NHP populations such as Vietnamese, Cambodian, and Cambodian/Indonesian mixed-origin cynomolgus macaques at Chinese breeding centers (Karl et al., 2017). In a similar study of pig-tailed macaques from multiple breeding centers, PacBio sequencing of MHC class I cDNA amplicons was used to define over 300 novel *Mane* sequences as well as 192 Mane-A and Mane-B regional haplotypes (Semler et al., 2018). Here, we have extended this PacBio circular consensus sequencing approach to full-length cDNA amplicons from MHC class II as well as class I transcripts.

In addition to PacBio sequencing, we made use of an extensive database of short tandem repeat (STR) data spanning the 5Mb MHC region that has been generated over multiple generations of this NIH-sponsored ICM breeding colony (Larsen et al., 2010). This is analogous to the approach that Otting *et al*. (Otting et al., 2012) employed to characterize extended MHC haplotypes in the BPRC cynomolgus macaque breeding groups. Doxiadis and colleagues (Doxiadis et al., 2013) employed a similar experimental strategy combining STR data with Sanger sequencing results in order to characterize the class I and class II transcript content of 176 founder haplotypes in a large breeding colony of rhesus macaques at the BPRC. Likewise, a related approach with pyrosequencing was utilized to characterize a closed colony of sooty mangabeys (*Cercocebus atys*) at the Yerkes National Primate Research Center for which STR data was available (Heimbruch et al., 2015). This study yielded 121 novel MHC class I transcript sequences that were assigned to 22 MHC haplotype that included combinations of both *Ceat-A* and *Ceat-B* transcripts. Here, we combine PacBio sequencing with MHC STR patterns to unambiguously assign specific *Mafa* class I and class II transcripts to the extended MHC haplotypes that are segregating in this pedigreed ICM population.

## Methods

### Animals

All animals evaluated in this study were members of the NIH-sponsored breeding colony at Alpha Genesis Inc. in Yemassee, SC, USA. that houses a cohort of approximately 350 ICM. This colony was established in 2002 and supplemented with additional groups of breeders in 2005 and 2010. The effective number of founding individuals was 79 for this breeding population. At least sixteen of these founders were born in Indonesia although the precise location of these births was unknown, and the remaining founders were reported by their distributors to be of Indonesian origin. Demographic information about the 295 individuals that were evaluated by sequencing in this study is provided in **Supplemental Figure 1**. Whole blood samples were obtained during routine health checks from this cohort for sequence analyses. These individuals spanned multiple generations which allowed us to use pedigree data to compare multiple offspring from common sires and dams. All animals were cared for according to the regulations and guidelines of the Institutional Care and Use Committee at Alpha Genesis Inc.

### STR haplotype patterns

STR analyses were completed as described previously (Penedo et al., 2005; Larsen et al., 2010) to determine parentage and place individuals within the pedigree. Next, STR allele sizes for a panel of 11 STR loci spanning the extended MHC region were compared between full-and half-siblings versus their parents in order to identify shared STR patterns that define the ancestral MHC haplotypes in this colony. Based on this analysis, each animal was assigned a pair of MHC haplotype codes, e.g. MHC-MAFA-NIAID1-00001 and MHC-MAFA-NIAID1-00002. **Supplemental Figure 2** provides a list of the STR allele patterns that are associated with each of the 100 extended haplotypes that were defined by PacBio sequencing in this study.

### RNA isolation, cDNA synthesis, and PCR amplification

RNA was isolated from whole blood in PAXgene Blood RNA tubes (Qiagen, Venlo, Netherlands) with a Maxwell 16 instrument using Maxwell 16 LEV SimplyRNA Blood kits according to the manufacturer’s protocol (Promega, Madison, WI, USA). Complementary DNA (cDNA) was then prepared using a SuperScript III First-Strand Synthesis System for RT-PCR by Invitrogen (Carlsbad, CA, USA). The conditions for this reaction were: 65°C for 5 min to anneal oligo dT primers; 4°C for 1 min, 50°C for 50 min, and 85°C for 5 min with Superscript enzyme; and finally, 37°C for 20 min after adding RNase H. These cDNA templates were amplified through 23-30 cycles of PCR using Phusion High-Fidelity master mix (New England Biolabs, Ipswich, MA, USA). For MHC class I sequencing, we used a cocktail of two forward and three reverse barcoded primers (**Supplemental Figure 3**). These 50 ul reactions included 1 ul cDNA, 0.1 uM primers, and Phusion High-Fidelity master mix. PCRs were run on an Applied Biosystems Thermal Cycler (ThermoFisher Scientific, Waltham, MA, USA) with reaction conditions of: 98°C for 3 min; 23 cycles of 98°C for 5 s, 60°C for 10 s, and 72°C for 20 s; followed by 72°C for 5 min. A similar process was used for MHC class II with a few distinctions (Karl et al., 2014). First, each of the class II loci required a separate PCR reaction and primers (**Supplemental Figure 3**). Additionally, we used 30 PCR cycles instead of 23, but with the same reaction conditions. All PCR products were monitored with FlashGels (Lonza, Basel, Switzerland) to confirm the anticipated length depending on locus. PCR amplicons were then subjected to two rounds of cleanup with the AMPure XP PCR purification kit (Agencourt Bioscience Corporation, Beverly, MA, USA) in a 0.7:1 ratio of beads to PCR sample. Purified amplicons were quantified using Qubit High Sensitivity kits (ThermoFisher Scientific, Waltham, MA, USA).

### PacBio library preparation and sequencing

For class I, amplicons were normalized to equal concentrations of 10 ng/ul in pools of 32-48 samples per PacBio library. For class II, we pooled 32-48 samples by loci, and each pool was quantified with Qubit High Sensitivity kits. These individual subpools were then combined into a final pool with 500ng DNA total in the ratio 3 DRB: 1 DQA: 1 DQB: 1 DPA: 1 DPB. Prior to sequencing, SMRTbell adaptors which form hairpins on the ends of the amplicons were ligated according to manufacturer’s protocol. Sequencing was completed on a PacBio Sequel instrument (PacBio, Menlo Park, CA, USA) at the University of Wisconsin-Madison Biotechnology Center.

### PacBio circular consensus sequence analysis

Raw sequence data was processed using the smrtlinkv6.0.0 command line toolset (https://www.pacb.com/support/software-downloads/). Briefly, smrtlink Dataset was run on raw sequencing data to produce a processed XML file. The resulting XML file was converted into BAM format containing circular consensus sequences (CCS) using the smrtlink CCS tool with the minimum and maximum length parameters set to 1000 and 1500 for MHC class I and 700 and 1000 for MHC class II. BAM files were then converted to fastq format using the smrtlink BAM2FASTQ tool. Fastq files were demultiplexed by barcodes using custom python code (https://github.com/dholab/Shortreed-et-al-Supplemental-Code). Long Amplicon Analysis (LAA) was run on each demultiplexed fastq file. The minimum and maximum length parameters were set to 1000 and 1500 for MHC class I, and 700 and 1000 for MHC class II. For both MHC class I and II, the Max_Clustering_Reads parameter was set to 5000 and Max_reads parameter was set to 20000.

Resulting amplicon sequences were mapped against the most recent library as of each sequencing run of known full-length *Mafa* MHC sequences in the NHP Immuno Polymorphism Database (IPD-MHC) using bbmap (http://sourceforge.net/projects/bbmap/) in semiperfect mode. Full-length LAA sequences that matched with perfect identity were removed from further analysis, as these represented known alleles. The remaining LAA sequences were mapped to a library of known partial *Mafa* MHC sequences using bbmap in semiperfect mode. Sequences that perfectly matched a partial sequence in the database were marked as putative extensions. Sequences that differed by one or more single nucleotide variants were marked as putative novel alleles. All of these putative novel and extended sequences were given temporary names and added to the database for the remainder of data processing. Next, reads for each animal were mapped against the database containing putative novels, putative extensions, and known sequences.

### DNA isolation, multiplex PCR and Illumina MiSeq analysis

Genomic DNAs were isolated from whole blood EDTA samples with a Maxwell 16 instrument using Maxwell 16 LEV Blood DNA kits according to the manufacturer’s protocol (Promega, Madison, WI, USA). These genomic DNAs were used as templates for PCR with a panel of primers that flank the highly polymorphic peptide binding domains encoded by exon 2 of class I (Mamu-A, -B, -I, -E) and class II (Mamu -DRB, -DQA, -DQB, -DPA and -DPB) loci. These PCR products were generated with a Fluidigm Access Array 48.48 (Fluidigm, San Francisco, CA, USA) which allows us to multiplex all reactions in a single experiment as described previously (Karl et al. 2014, 2017). After cleanup and pooling, these amplicons were sequenced on an Illumina MiSeq instrument (Illumina, San Diego, CA, USA). The resulting Illumina sequence reads were mapped against a custom reference database of *Mafa* class I and class II sequences as previously described (Caskey et al., 2019).

### Allele validation and haplotype analysis

Validation of novel and extended sequences was a multi-step process. First, we used Geneious (Biomatters, Auckland, New Zealand) software to visualize and confirm that the sequences had the expected start and stop amino acid sequences and that there were not any significant gaps or insertions such that the length was within the expected range for each locus. Next, we considered read support for a particular sequence relative to other reads in the same animal and compared across related animals. A potential novel allele that was supported by a minimum of three identical CCS reads in one or more animals was considered valid if there were not alleles of the same lineage with better read support within the same animal(s). These criteria prevent mischaracterization of PCR artifacts and sequencing errors as valid new MHC sequences.

*Mafa* class I and class II transcripts that are associated with extended MHC haplotypes in this breeding colony were assigned based on their identification in multiple individuals that share a specific MHC-MAFA-NIAID1 STR pattern. In a few cases where PacBio class I and/or class II sequencing results were only available for a single individual representing one of the less frequent MHC haplotypes, transcripts that remained after those associated with a more common, shared haplotype had been identified were inferred to segregate with these rare MHC haplotypes. The *Mafa-A, Mafa-B* and *Mafa-DRB* gene clusters are physically linked with a strong tendency to travel together as regional haplotypes (Karl et al., 2017). Since each of these gene clusters are separated by approximately 1Mb on chromosome 4 however, an occasional recombination event may occur that breaks the linkage between these haplotype blocks. To provide a concise description of each of these allele combinations, we developed a series of abbreviated Mafa-A, Mafa-B and Mafa-DRB haplotype designations (Karl et al., 2017). The specific *Mafa* transcript sequences that we have defined for each of these regional haplotypes in the Alpha Genesis breeding colony are summarized in **Supplemental Figures 4-6**.

## Results & Discussion

### Novel *Mafa* transcripts & extensions of partial *Mafa* sequences

In the initial phase of this study, we performed MHC class I allele discovery by PacBio circular consensus sequencing of full-length cDNA amplicons from a total of 295 ICM in this breeding colony. PacBio sequencing yielded an average of 1,668 MHC class I reads per animal that were either identical to known *Mafa* alleles, extensions of previously described partial *Mafa* sequences, or validated as novel *Mafa* allelic variants. These novel Mafa transcript sequences differed by one or more synonymous or nonsynonymous nucleotide changes compared to previously reported *Mafa* sequences available in IPD-MHC (Maccari et al., 2017). In total we identified 141 class I sequences in this study (Table 1); 90 of these transcripts represented novel sequences and there were 27 *Mafa-*A, 53 *Mafa-*B and 10 *Mafa-*I allelic variants. These new *Mafa* class I sequences have been deposited in the IPD-MHC Non-Human Primate Database (https://www.ebi.ac.uk/ipd/mhc/group/NHP) (de Groot et al., 2019).

**Table 1:**
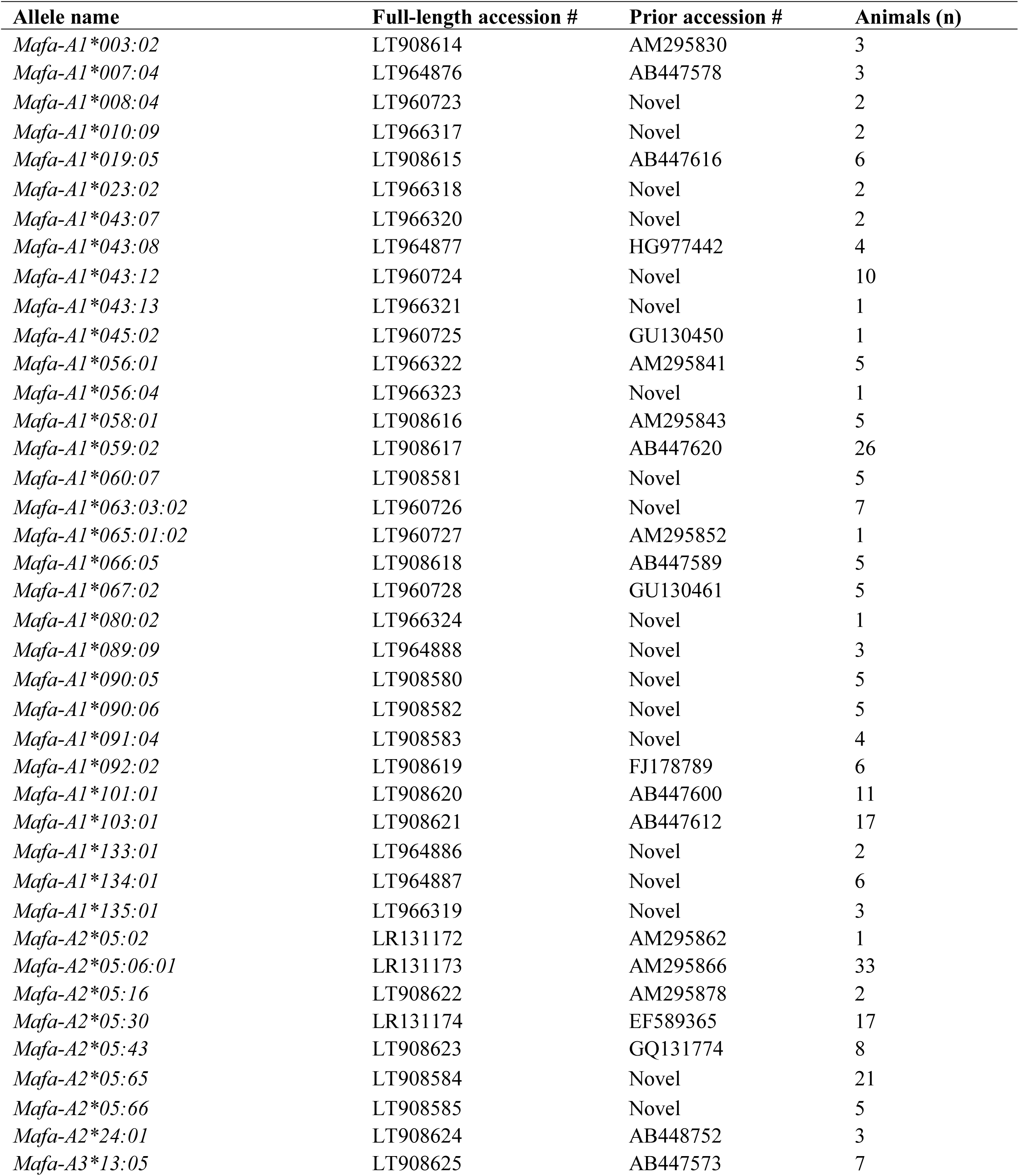
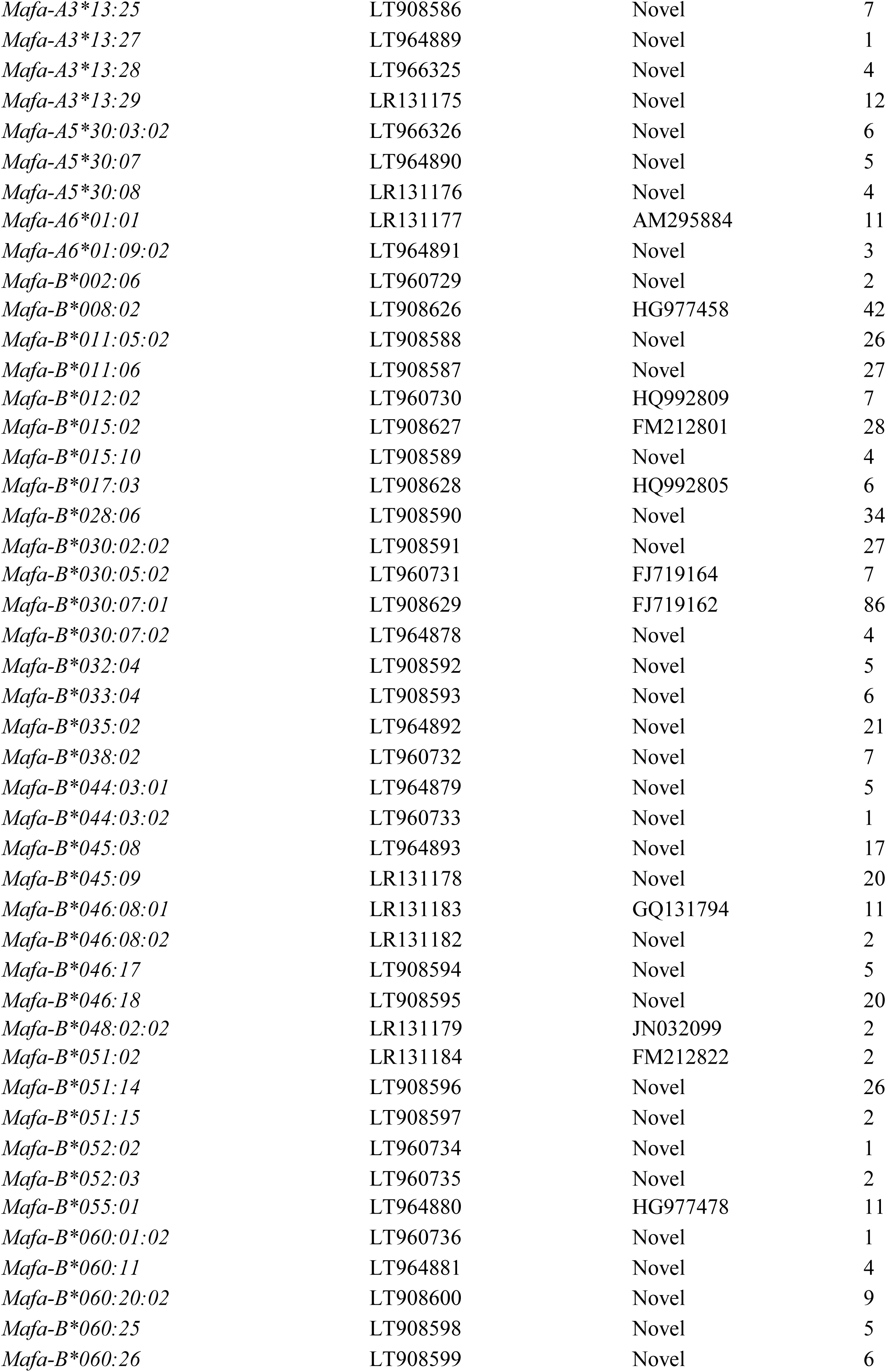
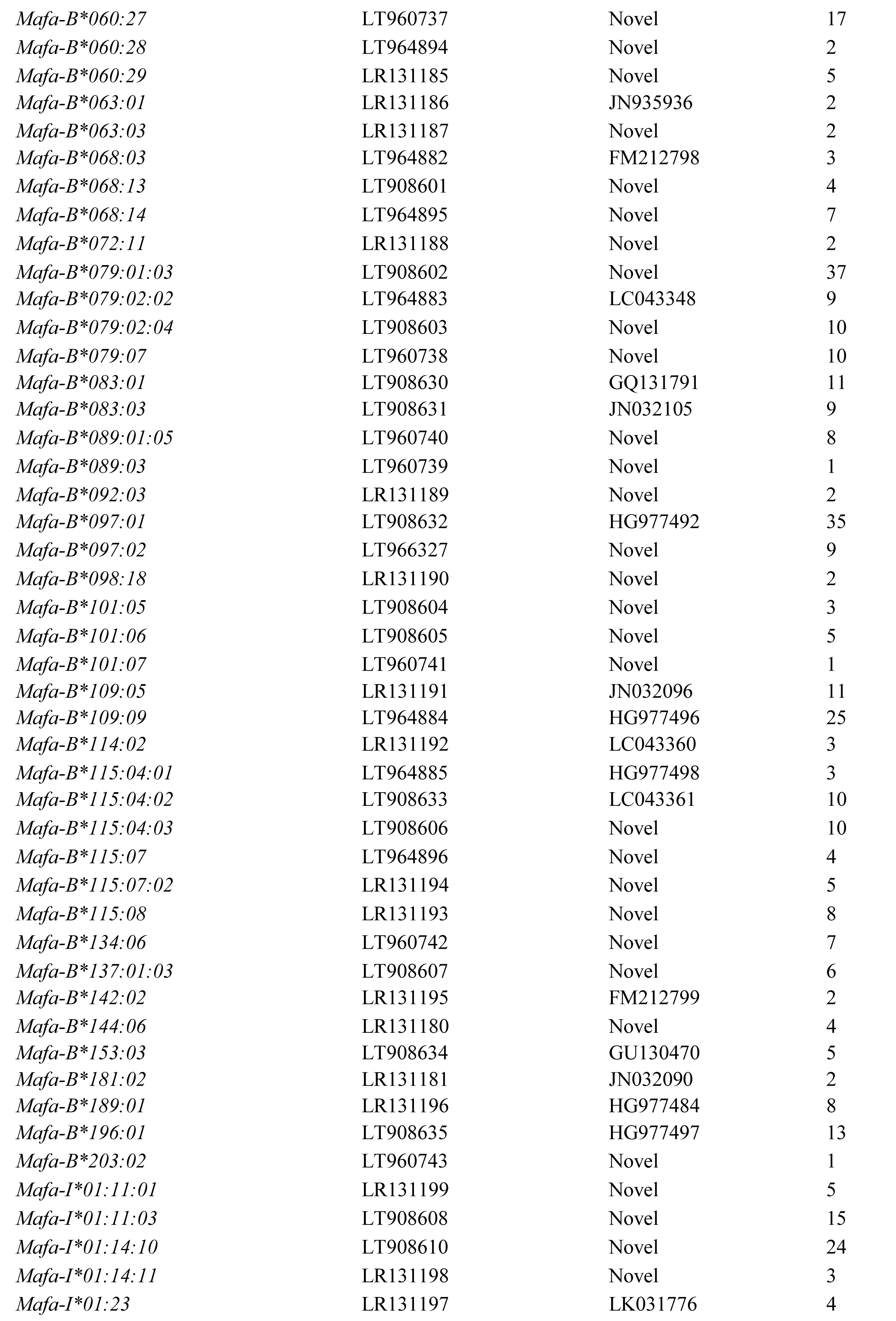
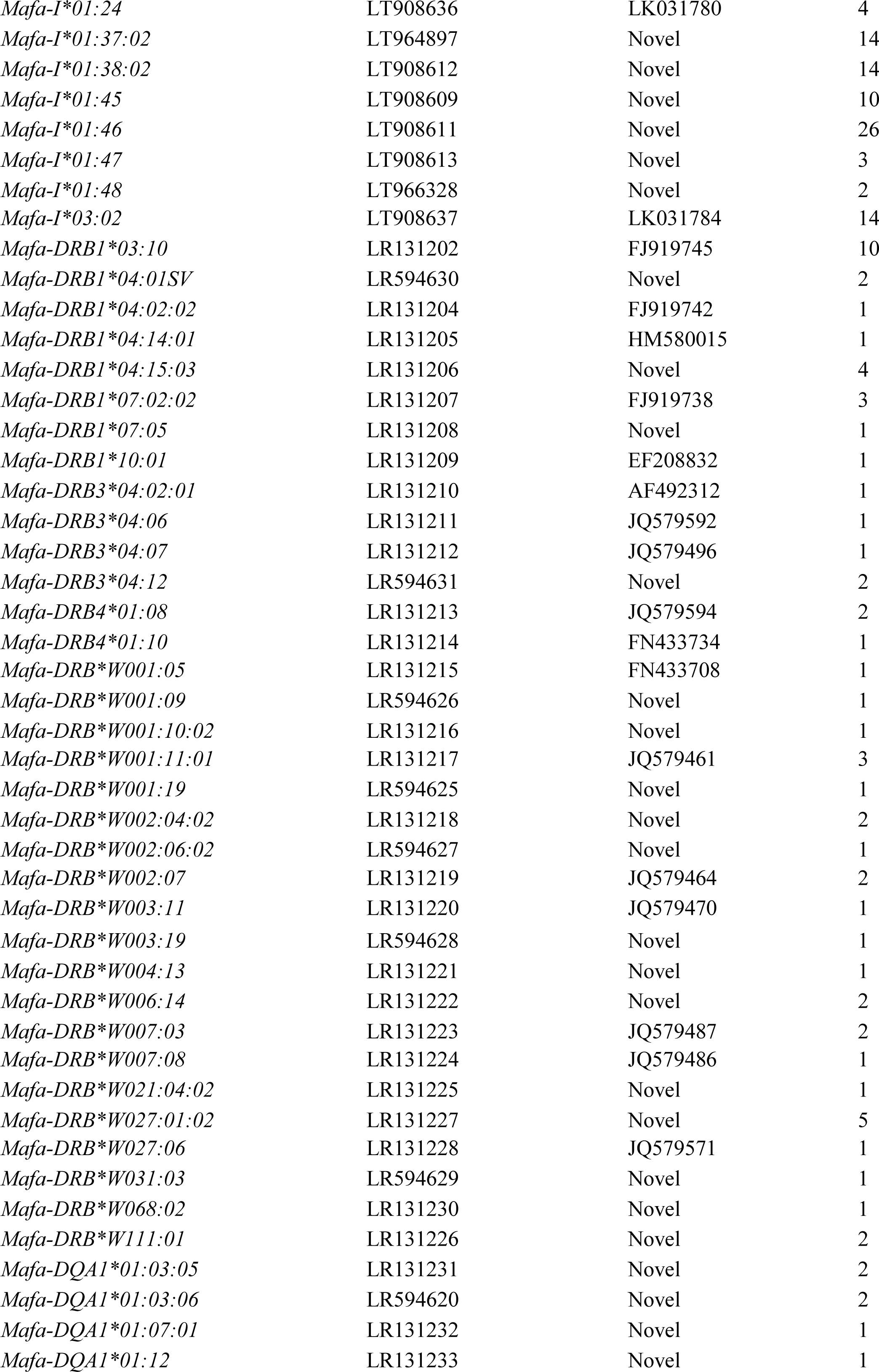
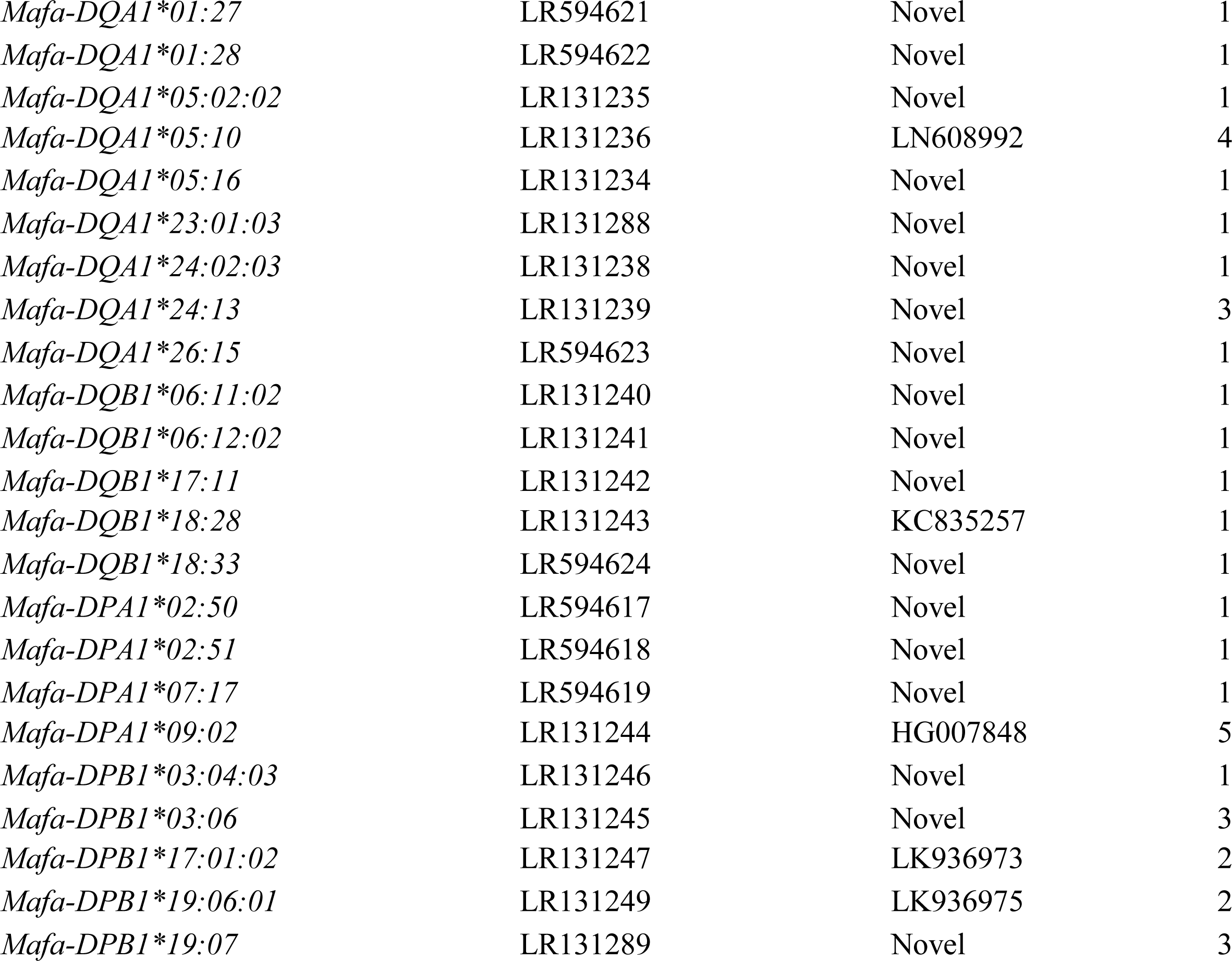
Summary of full-length ORF sequences identified in 295 animals. This table gives the official IPD-MHC allele nomenclature, GenBank accession number, and the total number of animals in which each sequence was observed. Extensions of previously described partial *Mafa* sequences also include accession numbers for the longest partial-length sequence previously described.

For characterization of MHC class II sequences, we adopted a targeted allele discovery strategy that made use of STR profiles for the MHC region that had been used to define extended MHC haplotypes in this ICM breeding colony. Based on knowledge of these STR patterns, we selected 76 representative animals that carried a total of 100 distinct ancestral MHC haplotypes. PacBio sequencing was performed with full-length cDNA amplicons for MHC class II loci and yielded a total of 3,857 MHC class II reads per animal. We identified a total of 61 additional full-length class II sequences. As summarized in Table 1, 39 of these class II sequences were novel and they included 17 *Mafa-*DRB, 12 *Mafa-*DQA1, 4 *Mafa-*DQB1, 3 *Mafa-*DPA1 and 3 *Mafa-*DPB1 allelic variants that have also been deposited in IPD-MHC (Table 1).

### Defining transcripts associated with regional and extended MHC haplotypes

After characterizing new class I and class II sequences with PacBio sequencing, we turned our efforts to defining the transcripts that are associated with the extended MHC haplotypes that are currently segregating in the Alpha Genesis breeding colony. Each of these extended MHC haplotypes have been defined by distinct patterns of allele sizes for a panel of eleven STR markers that span the 5Mb MHC genomic region (**Supplemental Figure 2**) (Penedo et al., 2005; Larsen et al., 2010). As illustrated in Figure 1, each extended MHC haplotype in cynomolgus macaques encodes multiple *Mafa-*A*, Mafa-*B *and Mafa-*DRB transcripts. In addition, there is considerable copy number variation between haplotypes as well as a wide range of steady-state RNA levels in whole blood for *Mafa-*A and *Mafa-*B transcripts from different loci. In order to describe the complex, high-resolution *Mafa* genotypes associated with each of the extended MHC haplotypes, we developed an abbreviated, regional haplotype nomenclature system for the Mafa-A, Mafa-B and Mafa-DRB gene clusters that is similar to MHC allele nomenclature (Figure 1). Each of these regional haplotype designations begins with the species name (Mafa) and a “diagnostic” allele lineage for a major transcript that is expressed at high steady-state levels, e.g. A018. Regional haplotypes that differ in their combinations of major transcripts are distinguished by a “.” delimiter followed by two digits in ascending order of their identification in various cynomolgus macaque populations. A final “.” delimiter separates closely related haplotypes that share the same major transcript lineages but differ due to nonsynonymous and/or synonymous allelic variants. For example, the regional Mafa-A018.01.05 haplotype illustrated in Figure 1 includes major *Mafa-A1*018:05:02* and minor *Mafa-A2*05:66* transcripts and it represents the fifth combination of allelic variants noted to date. Using this abbreviated haplotype system, the extended MHC haplotypes defined by the MHC-MAFA-NIAID1-000061 STR pattern can be described by the following string: A018.01.05|B147.04.02|DRBW020.04.02|DQA26_01_01|DQB15_01_02|DPA02_13_02|DPB15_02.

**Figure 1:**
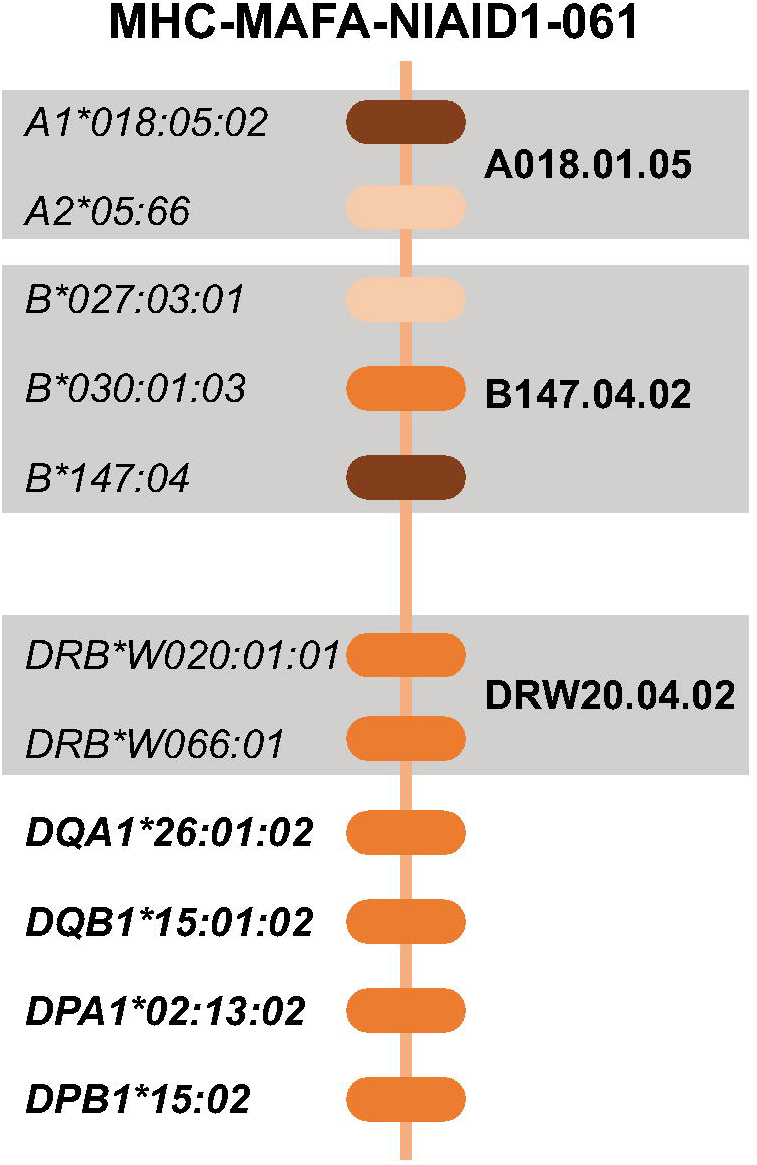
An example of an extended MHC haplotype defined by the MHC-MAFA-NIAID1-000061 STR pattern. For *Mafa*-*A*, -*B*, and *-DR*B, multiple transcripts with a wide range of steady-state RNA levels as indicated by the intensity of the color of the locus bubble, together make up the regional haplotype in bold. For the remaining class II loci, each gene is indicated by a single transcript in bold font.

Genotyping results for a representative group of four progeny from sire CX7K is presented in Figure 2 to illustrate how PacBio sequencing data is used to define the *Mafa* transcripts that are associated with extended MHC haplotypes. In this example, each of the offspring inherited the same paternal MHC haplotype that is defined by the MHC-MAFA-NIAID1-000048 STR pattern. The values in the body of Figure 2 indicate the PacBio sequence read support for each of the class I and class II transcripts that were identified in each animal and color coding denotes the haplotype for each sequence. Transcripts highlighted in yellow are shared by all four of these offspring and thus are assigned to the paternal haplotype that was inherited from the CX7K sire. The *Mafa-B* gene cluster of this extended haplotype includes a diagnostic major *Mafa-B*028:06* transcript as well as six additional *Mafa*-B transcripts with a range of steady-state RNA levels. To simplify communicating these genotyping results we abbreviate this combination of Mafa-B loci as the Mafa-B028.07.01 regional haplotype. In the same way, the transcripts that are unique in each of these progeny are inferred to be associated with their maternal haplotypes (Figure 2). For instance, the combination of *Mafa-B*147:04, Mafa-B*030:01:03* and *Mafa-B*027:03:01* transcripts expressed by H592 are designated as the Mafa-B147.04.02 regional haplotype. This Mafa-B haplotype is one building block of the MHC-MAFA-NIAID1-000061 extended haplotype that was inherited from dam CX95.

**Figure 2:**
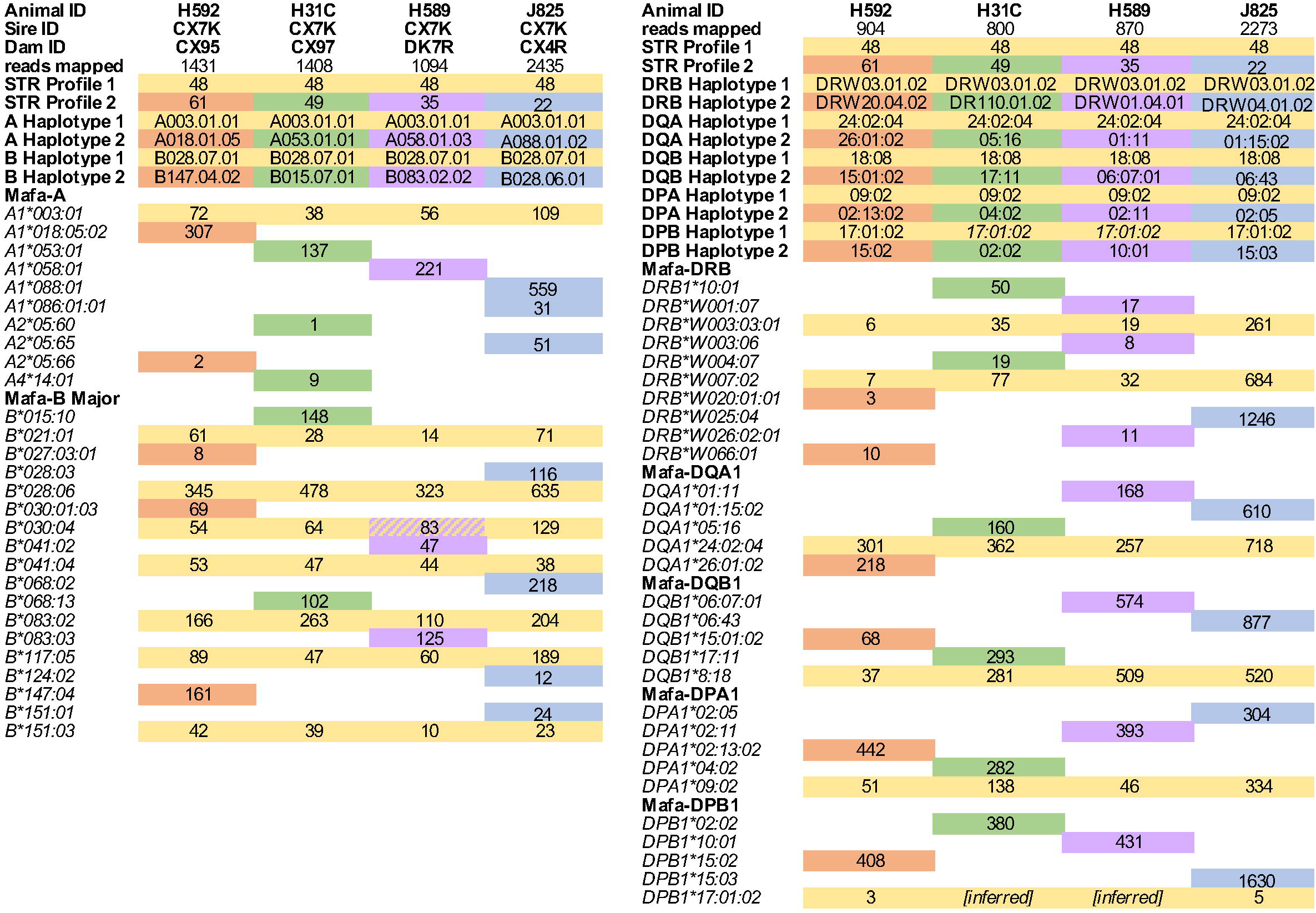
PacBio sequencing results for four of the 295 macaques in this study. MHC class I results are shown on the left and class II data is on the right side. For each macaque, we list the animal ID, sire ID, dam ID, and number reads mapped with PacBio sequencing. Each of these animals shares a common sire and inherited the paternal haplotype containing the MHC-MAFA-NIAID1-000048 STR pattern (yellow). Each of the progeny express a series of class I and class II transcripts in common (yellow) that are represented by the number of identical PacBio sequences reads that were detected in each animal. These co-segregating transcripts are used to define the abbreviated regional haplotypes listed below the STR profiles. The remaining transcripts expressed by each animal are highlighted in a different color and inferred to segregate as the maternal haplotype inherited from their respective dams. Diagonally shaded boxes indicate that a transcript such as *Mafa-B*030:04* is included on both the paternal and maternal extended MHC haplotypes. If PacBio read support is low, but a sequence is observed in related animals and/or MiSeq data, it is noted as inferred, *e.g.*, *Mafa-DPA1*17:01:02* for the extended haplotype defined by the shared MHC-MAFA-NIAID1-000048 STR pattern.

By iterating this process with PacBio genotyping results from groups of related animals that share STR patterns from common ancestors, we defined the class I and class II transcripts that are associated with 100 extended MHC haplotypes in this breeding colony as summarized in Table 2. **Supplemental Figures 4 – 6** provide lists of the *Mafa-A*, *Mafa-B* and *Mafa-DRB* transcripts that are associated with each of the abbreviated, regional Mafa haplotype designations displayed in Table 2. Complete PacBio read support per transcript is available for all animals sequenced in this study in **Supplemental Figure 7**. In a few cases where PacBio read support was low, these results were augmented with lineage-level MiSeq genotyping results with exon 2 genomic DNA amplicons as shown in **Supplemental Figure 8** (Karl et al., 2017).

**Table 2:** Mafa haplotype definitions for 100 extended MHC haplotypes defined by MHC-MAFA-NIAID1 STR patterns. STR allele sizes for each extended MHC haplotype are listed in Supplemental Figure 2. The number of each haplotype observed (# Obs) was inferred based on animals with STR patterns matching the 100 extended MHC haplotypes characterized in this study (n = 1146). Mafa class I and class II transcripts associated with each of the abbreviated Mafa-A, Mafa-B and Mafa-DRB regional haplotypes are listed in Supplemental Figures 4 - 6. Individual Mafa-DQA1, DQB1, DPA1 and DPB1 alleles associated with each extended MHC haplotype are listed in the final 4 columns.

| MHC-MAFA-NIAID1- | # Obs. | Mafa-A | Mafa-B | Mafa-DRB | Mafa-DQA | Mafa-DQB | Mafa-DPA | Mafa-DPB |
| --- | --- | --- | --- | --- | --- | --- | --- | --- |
| 000005 | 125 | A007.01.03 | B008.02.02 | DRW33.01.01 | 24:02:01 | 18:08 | 02:13:02 | 15:02 |
| 000002 | 62 | A010.01.05 | B008.02.01 | DR104.06.03 | 01:03:05 | 06:12 | 02:13:02 | 15:02 |
| 000008 | 45 | A060.02.01 | B015.05.01 | DR103.02.02 | 01:24:01 | 06:13:01 | 04:02 | 03:04:01 |
| 000001 | 35 | A018.01.03 | B018.02.01 | DR103.01.02 | 24:05 | 18:05 | 02:11 | 10:01 |
| 000010 | 35 | A022.01.06 | B147.03.01 | DR103.06.01 | 01:03:02 | 06:18 | 07:07 | 19:06:01 |
| 000025 | 34 | A054.01.01 | B013.08.01 | DR401.02.01 | 01:21 | 06:24:02 | 04:02 | 03:04:01 |
| 000007 | 30 | A103.01.01 | B015.01.02 | DR103.20.01 | 01:03:02 | 06:18 | 04:02 | 03:04:01 |
| 000006 | 29 | A004.01.02 | B045.02.02 | DR103.12.01 | 01:07:01 | 06:14 | 04:02 | 03:06 |
| 000048 | 29 | A003.01.01 | B028.07.01 | DRW03.01.02 | 24:02:04 | 18:08 | 09:02 | 17:01:02 |
| 000009 | 28 | A008.01.02 | B004.03.01 | DRW21.01.01 | 05:10 | 17:02:01 | 02:14:01 | 01:05 |
| 000022 | 26 | A088.01.02 | B028.06.01 | DRW04.02.01 | 01:15:02 | 06:43 | 02:05 | 15:03 |
| 000028 | 26 | A010.01.05 | B028.05.01 | DR103.18.01 | 24:05 | 18:05 | 02:13:02 | 15:02 |
| 000132 | 25 | A089.01.03 | B056.04.01 | DR110.04.01 | 05:03:01 | 16:01 | 02:05 | 15:03 |
| 000018 | 23 | A043.01.04 | B076.02.01 | DRW04.01.01 | 01:12 | 06:14 | 07:04:01 | 21:02 |
| 000003 | 22 | A038.01.03 | B024.01.01 | DR103.06.01 | 01:03:02 | 06:18 | 07:01 | 21:02 |
| 000019 | 21 | A007.01.03 | B002.01.02 | DR103.05.01 | 05:01:01 | 17:03 | 02:18 | 06:06:02 |
| 000072 | 20 | A019.01.02 | B090.03.01 | DRW33.01.01 | 24:04 | 18:05 | 07:02 | 19:03 |
| 000013 | 18 | A043.01.05 | B004.01.01 | DR103.06.01 | 01:03:05 | 06:12 | 02:13:02 | 15:02 |
| 000035 | 17 | A058.01.03 | B083.02.02 | DRW01.04.01 | 01:11 | 06:07:01 | 02:15:01 | 10:01 |
| 000037 | 15 | A068.01.01 | B008.02.01 | DRW05.01.01 | 24:03 | 18:01:02 | 02:27 | 01:01:01 |
| 000117 | 15 | A019.01.01 | B028.01.04 | DR104.13.01 | 24:07 | 18:28 | 02:01 | 25:03 |
| 000036 | 14 | A019.01.01 | B052.02.01 | DRW01.02.03 | 23:01:01 | 18:31 | 04:02 | 03:04:01 |
| 000058 | 14 | A010.01.03 | B056.04.02 | DRW04.01.01 | 01:12 | 06:14 | 02:15:01 | 10:01 |
| 000063 | 14 | A010.01.05 | B017.02.01 | DR103.05.01 | 05:01:01 | 17:03 | 02:09:02 | 15:09 |
| 000004 | 13 | A101.01.01 | B016.01.02 | DRW04.01.01 | 01:12 | 06:08 | 02:15:01 | 10:01 |
| 000031 | 12 | A054.01.02 | B069.07.01 | DRW04.01.01 | 01:12 | 06:14 | 02:09:02 | 15:09 |
| 000043 | 12 | A088.01.01 | B184.01.03 | DRW01.04.01 | 01:11 | 06:07:01 | 02:26 | 25:03 |
| 000032 | 11 | A019.02.01 | B090.01.01 | DRW21.02.01 | 24:02:04 | 18:04 | 11:01 | 16:01 |
| 000061 | 11 | A018.01.05 | B147.04.02 | DRW20.04.02 | 26:01:02 | 15:01:02 | 02:13:02 | 15:02 |
| 000064 | 11 | A070.01.01 | B181.02.01 | DR103.06.01 | 01:03:02 | 06:18 | 07:07 | 19:06:01 |
| 000130 | 11 | A060.01.03 | B101.02.01 | DR103.06.01 | 01:03:02 | 06:18 | 02:13:02 | 15:02 |
| 000052 | 10 | A053.01.01 | B004.01.02 | DRW21.02.01 | 23:01:01 | 18:04 | 07:03 | 19:06:02 |
| 000076 | 11 | A135.01.01 | B008.02.03 | DRW01.04.01 | 01:11 | 06:07:01 | 13:03 | 04:01 |
| 000077 | 10 | A066.01.04 | B017.03.01 | DR104.05.02 | 01:06:01 | 06:11:02 | 09:02 | 17:01:02 |
| 000011 | 9 | A076.01.01 | B028.06.03 | DR107.02.01 | 24:05 | 18:05 | 04:02 | 03:04:03 |
| 000024 | 9 | A043.01.07 | B045.02.01 | DR103.06.01 | 01:03:02 | 06:18 | 02:15:01 | 10:01 |
| 000027 | 9 | A091.01.01 | B147.02.01 | DRW03.01.03 | 24:02:03 | 18:08 | 02:26 | 25:04 |
| 000030 | 9 | A070.01.01 | B013.08.01 | DRW21.02.01 | 23:01:01 | 18:04 | 09:02 | 17:01:02 |
| 000047 | 9 | A053.01.01 | B004.01.02 | DRW01.02.03 | 24:03 | 18:04 | 04:02 | 03:04:01 |
| 000097 | 9 | A018.01.04 | B004.01.01 | DRW01.03.01 | 05:01:01 | 17:03 | 02:15:01 | 10:01 |
| 000131 | 9 | A063.01.04 | B104.01.02 | DRW03.01.02 | 24:02:03 | 18:08 | 13g | 04:01 |
| 000016 | 8 | A101.02.01 | B004.01.03 | DR107.03.01 | 24:13 | 18:06 | 02:13:02 | 15:02 |
| 000020 | 8 | A041.01.02 | B047.01.02 | DR104.10.04 | 26:06 | 17:07:01 | 02:16 | 06:08 |
| 000029 | 8 | A018.01.07 | B083.02.02 | DR104.08.02 | 01:09:01 | 06:01:02 | 11:01 | 16:01 |
| 000078 | 8 | A092.02.02 | B056.04.02 | DR107.04.01 | 01:11 | 06:07:01 | 07:01 | 21:02 |
| 000079 | 8 | A041.01.02 | B047.01.02 | DR503.01.01 | 05:02:02 | 17:06:02 | 02:14:01 | 01:05 |
| 000053 | 7 | A003.01.01 | B194.01.01 | DR103.19.01 | 01:04 | 06:01:01 | 02:06:02 | 07:01 |
| 000081 | 7 | A088.01.01 | B056.07.01 | DR304.09.01 | 01:03:06 | 06:12:02 | 02:14:01 | 01:05 |
| 000023 | 6 | A010.01.05 | B013.09.01 | DR110.03.01 | 01:24:02 | 06:13:01 | 02:13:02 | 15:02 |
| 000044 | 6 | A067.01.03 | B090.02.02 | DR103.19.01 | 01:07:01 | 06:14 | 11:01 | 16:01 |
| 000050 | 6 | A006.01.01 | B013.08.01 | DRW33.01.01 | 24:04 | 18:05 | 13g | 04:01 |
| 000067 | 6 | A067.01.01 | B048.02.01 | DR104.06.03 | 01:03:05 | 06:12 | 02:13:02 | 15:02 |
| 000082 | 6 | A089.01.02 | B015.06.01 | DRW21.01.01 | 05:10 | 17:02:01 | 07:07 | 19:07 |
| 000085 | 6 | A077.01.01 | B101.03.01 | DRW01.05.01 | 01:13 | 06:17:02 | 02:14:01 | 01:05 |
| 000091 | 6 | A088.01.03 | B151.02.01 | DR103.30.01 | 01:27 | 06:18 | 13:02 | 06:06:01 |
| 000033 | 5 | A097.01.02 | B002.03.01 | DRW21.01.01 | 05:10 | 17:02:01 | 02:15:01 | 10:01 |
| 000034 | 5 | A010.03.01 | B056.06.03 | DRW04.01.01 | 01:12 | 06:14 | 02:09:02 | 15:09 |
| 000049 | 5 | A053.01.01 | B015.07.01 | DR110.01.02 | 05:16 | 17:11 | 04:02 | 02:02 |
| 000065 | 5 | A070.01.01 | B101.03.01 | DR110.04.01 | 05:03:01 | 16:01 | 13:01 | 09:02 |
| 000083 | 5 | A010.01.06 | B013.08.01 | DRW01.04.02 | 01:11 | 06:07:01 | 02:16 | 06:08 |
| 000086 | 5 | A024.02.01 | B104.02.04 | DRW04.01.01 | 01:12 | 06:14 | 02:15:01 | 10:01 |
| 000109 | 5 | A008.01.03 | B101.03.02 | DR103.02.02 | 01:24:02 | 06:13:01 | 07:07 | 19:06:01 |
| 000134 | 5 | A010.01.05 | B013.08.01 | DR103.12.01 | 01:07:01 | 06:14 | 07:04:01 | 20:01 |
| 000135 | 5 | A010.01.05 | B052.02.02 | DR304.08.01 | 01:28 | 06:20 | 02:51 | 10:01 |
| 000136 | 5 | A070.01.01 | B076.01.02 | DR304.09.01 | 01:03:06 | 06:12:02 | 02:11 | 10:01 |
| 000015 | 4 | A079.01.01 | B004.01.01 | DRW01.02.02 | 24:11 | 18:13 | 04:02 | 03:04:01 |
| 000040 | 4 | A067.01.02 | B012.01.01 | DR107.03.01 | 24:13 | 18:06 | 07:04:01 | 21:01 |
| 000051 | 4 | A043.01.08 | B017.02.02 | DRW01.03.01 | 01:11 | 06:07:01 | 13:03 | 04:01 |
| 000056 | 4 | A063.01.02 | B075.01.01 | DR110.04.01 | 05:03:01 | 16:01 | 13:01 | 09:02 |
| 000057 | 4 | A007.01.04 | B012.01.01 | DR103.06.01 | 01:03:02 | 06:18 | 07:07 | 19:06:01 |
| 000059 | 4 | A056.03.01 | B004.02.01 | DRW04.01.01 | 01:12 | 06:14 | 07:04:01 | 20:01 |
| 000060 | 4 | A067.01.03 | B147.04.01 | DRW01.02.03 | 23:01:01 | 18:31 | 02:27 | 01:01:01 |
| 000087 | 4 | A018.01.06 | B099.01.01 | DRW03.01.02 | 24:02:03 | 18:08 | 02:26 | 15:06 |
| 000090 | 4 | A010.01.04 | B013.08.01 | DR103.19.01 | 01:07:01 | 06:14 | 13:03 | 04:01 |
| 000098 | 4 | A004.01.02 | B090.02.01 | DRW07.01.01 | 23:01:03 | 18:31 | 02:15:01 | 15:02 |
| 000102 | 4 | A004.01.02 | B056.06.02 | DRW01.04.01 | 01:11 | 06:07:01 | 02:16 | 06:08 |
| 000104 | 4 | A088.01.01 | B184.01.02 | DRW04.01.01 | 01:12 | 06:14 | 02:15:01 | 10:01 |
| 000107 | 4 | A043.01.06 | B028.06.02 | DR304.10.01 | 26:15 | 18g | 07:01 | 21:02 |
| 000115 | 4 | A033.01.03 | B045.01.03 | DR104.12.01 | 01:06:01 | 06:11 | 13:03 | 04:01 |
| 000128 | 4 | A004.01.02 | B056.06.03 | DR104.08.02 | 01:09:01 | 06:01:02 | 11:01 | 16:01 |
| 000045 | 3 | A076.01.01 | B013.08.01 | DR104.12.01 | 01:06:01 | 06:11 | 02:27 | 01:01:01 |
| 000062 | 3 | A066.01.02 | B203.01.01 | DR107.04.02 | 01:11 | 06:07:01 | 02:14:01 | 01:05 |
| 000068 | 3 | A019.02.01 | B090.01.01 | DRW21.02.01 | 23:01:01 | 18:04 | 11:01 | 15:08 |
| 000075 | 3 | A070.01.01 | B056.06.01 | DRW01.04 | 01g1 | 06g3 | 02g7 | DPB-unk |
| 000105 | 3 | A079.01.01 | B076.01.01 | DR103.21.01 | 24g1 | 18:08 | 04:02 | 03g |
| 000118 | 3 | A101.01.01 | B101.04.01 | DRW21.01.01 | 05:10 | 17:02:01 | 07:07 | 19:07 |
| 000119 | 3 | A067.01.01 | B013.10.01 | DR107.04.01 | 01:11 | 06:07:01 | 04:02 | 03:04:02 |
| 000121 | 3 | A023.01.01 | B018.02.02 | DR103.05.01 | 05:01:01 | 17:03 | 02:11 | 10:01 |
| 000126 | 3 | A088.01.01 | B045.02.03 | DRW01.04.01 | 01:11 | 06:07:01 | 02:11 | 10:01 |
| 000012 | 2 | A066.01.05 | B083.05.01 | DRW04.03.01 | 01:03:02 | 06:36 | 02:15:01 | 10:01 |
| 000092 | 2 | A080.01.01 | B083.02.02 | DR107.03.01 | 24:13 | 18:06 | 07:04:01 | 15:02 |
| 000111 | 2 | A032.02.02 | B015.07.01 | DR503.01.01 | 24:11 | 18g1 | 07:03 | 19g |
| 000129 | 2 | A056.03.02 | B012.01.01 | DRW01.02.03 | 23:01:01 | 18:31 | 02:50 | 01:01:01 |
| 000133 | 2 | A092.01.03 | B101.04.01 | DR103.20.01 | 24:04 | 18:05 | 04:02 | 03:04:01 |
| 000026 | 1 | A006.01.02 | B002.04.01 | DRW21.01.01 | 05:16 | 17:02:01 | 07:07 | 03:06 |
| 000039 | 1 | A045.01.03 | B052.01.01 | DR107.02.01 | 24:05 | 18:05 | 07:02 | 19:03 |
| 000116 | 1 | A062.01.01 | B008.03.01 | DR103.01.02 | DQA-unk | 18:01:01 | 09:02 | 10:01 |
| 000127 | 1 | A004.01.02 | B069.08.01 | DRW01.06.01 | 01:03:02 | 06:20 | DPA-unk | 04:01 |
| 000137 | 1 | A004.01.02 | B083.05.02 | DRW21.02.01 | 23:01:01 | 18:04 | 09:02 | 17:01:02 |
| 000138 | 1 | A018.01.03 | B018.02.01 | DRW33.01 | 24g1 | 18:08 | 02g3 | 15g3 |

As a result of these efforts, we identified 70 distinct combinations of *Mafa-A* transcripts (**Supplemental Figure 4**) and 78 distinct combinations of *Mafa-B* transcripts (**Supplemental Figure 5**). At least 45 combinations of *Mafa-DRB* transcripts were also observed in this cohort (**Supplemental Figure 6**). Since *Mafa-DQ* and *Mafa-DP* haplotypes each only include single genes for the alpha and beta chains, the individual alleles for these four loci are listed for each extended MHC haplotype in Table 2. These results indicate that there are 45 distinct combinations of *Mafa-DQA1/ Mafa-DQB1* alleles and 38 different pairs of *Mafa-DPA1/ Mafa-DPB1* alleles. Many of these Mafa-DQ and Mafa-DP haplotypes confirm previous observations for individuals from this and other breeding colonies (Otting et al., 2017).

The majority of the regional Mafa-A, Mafa-B and Mafa-DRB haplotypes detected in this breeding colony are restricted to one or two extended MHC haplotypes (**Supplemental Figures 4 – 6**). This observation is consistent with a previous study by Otting and colleagues that characterized 32 multi-locus MHC haplotypes for pedigreed cynomolgus macaque breeding groups at the Biomedical Primate Research Center (BPRC) in the Netherlands (Otting et al., 2012). Several Mafa-A and Mafa-B regional haplotypes are shared by multiple extended haplotypes however and do appear to be somewhat enriched in the Alpha Genesis breeding colony. For example, the Mafa-A004.01.02, Mafa-A010.01.05 and Mafa-B013.08.01 haplotypes are each associated with six extended MHC haplotypes (**Supplemental Figures 4 and 5**). These extended MHC haplotypes do no not appear to result from simple recombination events in the class I region that arose in recent ancestors of this breeding colony. As illustrated in Figure 3, the STR patterns flanking the Mafa-A004.01.02 haplotypes provide evidence of a conserved genomic region that is marked by three to six common telomeric and centromeric STR alleles in four extended MHC haplotypes, but each of these Mafa-A004.01.02 haplotype blocks are linked to distinct Mafa-B and class II haplotypes. The first three STRs (222I18, MOG-CA and 151L13) reside approximately 03.-0.6 Mb telomeric to the *Mafa-A* gene cluster and the next three STRs (162B17A, 162B17B and 246K06) are localized a comparable distance from the centromere (Daza-Vamenta et al., 2004; Kean et al., 2012) where they flank the *Mafa-E* genes. Interestingly, the major *Mafa-A1*004:03* transcript that is included as part of this Mafa-A004.01.02 haplotype was originally identified as the most common *Mafa-A1* sequence identified in a cohort of ICM (Kita et al., 2009). In contrast, the STR patterns flanking the regional Mafa-A010.01.05 haplotypes are essentially completely distinct (Figure 3), suggesting that these six extended MHC haplotypes are truly ancient configurations. Unlike the extended cynomolgus macaque MHC haplotypes described here, the Indian-origin rhesus macaque population is characterized by a relatively limited number of Mamu-A, Mamu-B and Mamu-DRB regional haplotype blocks that have been shuffled by recombination to generate extended MHC haplotype diversity (Doxiadis et al., 2013; Karl et al., 2013; Wiseman et al., 2013).

**Figure 3:**
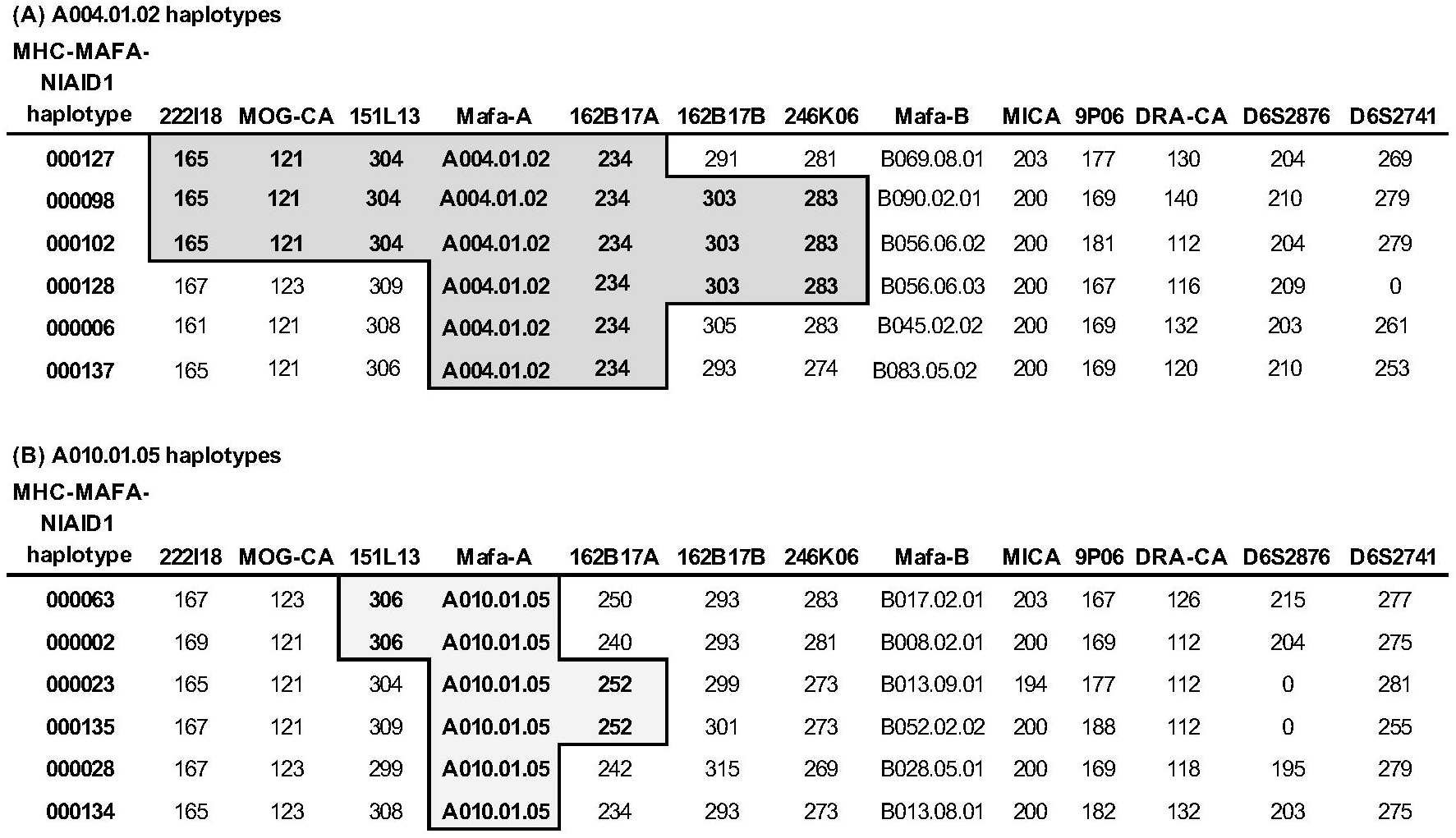
STR patterns for extended MHC haplotypes that share Mafa-A004.01.02 or Mafa-A010.01.05 haplotypes. STR loci are displayed from telomere (left) to centromere (right) and STR allele sizes that define each extended haplotype are listed. Shading and bold face highlights the STR alleles that are shared between multiple extended haplotypes that include Mafa-A004.01.02 (**A**) or Mafa-A010.01.05 (**B**) haplotypes.

The availability of phased STR patterns for a significant subset of the macaques in this pedigreed breeding colony allowed us to infer MHC genotypes based on haplotype definitions listed in Table 2 and **Supplemental Figures 4-6** for nearly twice as many individuals as those who were sequenced directly. In this study, we characterized the Mafa class I and class II transcripts that are associated with 100 of the most common extended MHC haplotypes that are currently segregating in this breeding colony. These 100 extended haplotypes account for 96.1% (1146 of 1192) of the total MHC haplotypes defined by STR results that are available for this breeding colony (Table 2). Distinct STR patterns have been identified for an additional 27 ancestral MHC haplotypes in this colony, but these remaining haplotypes were only detected in one to three individuals each. These rare MHC haplotypes were only detected in a total of 46 individuals and therefore only account for 3.9% of all MHC haplotypes for animals where STR data are available.

Overall the distribution of extended MHC haplotypes in this breeding colony was relatively uniform for animals whose STR patterns were known (Table 2). The extended MHC haplotype defined by the MHC-MAFA-NIAID1-000005 STR pattern was exceptional in this regard since it was observed in 121 individuals, four of whom were homozygous. The abundance of the MHC-MAFA-NIAID1-000005 haplotype reflects the fact that three of six founder males (CM94, CM9P and CX7K) who initiated breeding in this colony between 2002 and 2005 each carried this haplotype. The available pedigree records only show that each of these founder males had different dams but their sires were unknown so they could be paternal half-siblings and at the very least must share a recent common ancestor. Together these three males sired a total of 122 progeny at Alpha Genesis for whom STR results are available and 59 of these offspring inherited the MHC-MAFA-NIAID1-000005 haplotype. In contrast, the next most common extended MHC haplotype, MHC-MAFA-NIAID1-000002, was introduced in this colony by a single founder male (CM51). This haplotype has been observed in 62 individuals (Table 2), 25 of whom were sired by CM51. As expected, the remaining pair of founder sires (CH2D and 061973) also each introduced extended MHC haplotypes (MHC-MAFA-NIAID1-000022, MHC-MAFA-NIAID1-000028, MHC-MAFA-NIAID1-000132 and MHC-MAFA-NIAID1-000013) that are also among those most frequently observed in this colony (Table 2).

Interestingly, the most common extended MHC haplotypes described above (MHC-MAFA-NIAID1-000005 and MHC-MAFA-NIAID1-000002) are both characterized by major *Mafa-B*008* transcripts that are closely related allelic variants. The associated Mafa-B008.02.02 and Mafa-B008.02.01 regional haplotypes are distinguished by *Mafa-B*008:02* and *Mafa-B*195:01* versus *Mafa-B*008:03* and *Mafa-B*195:02* major transcripts, respectively (**Supplemental Figure 5**). Fortunately, the rhesus macaque orthologue (*Mamu-B*008:01*) of these *Mafa-B*008* sequences encodes one of the most thoroughly characterized MHC class I proteins in NHPs due to its strong association with exceptional control of simian immunodeficiency virus replication (Loffredo et al., 2007). Loffredo and coworkers have performed extensive epitope mapping studies and defined a detailed peptide binding motif for the Mamu-B*008:01 protein (Loffredo et al., 2009). Since the predicted protein product of Mafa-B*008:02 only differs from its rhesus counterpart by a single V189M substitution in the alpha 2 domain that is not expected to be involved in peptide binding, it is expected to bind the same spectrum of peptides as those described for Mamu-B*008:01. The predicted Mafa-B*008:03 protein (associated with the MHC-MAFA-NIAID1-000002 haplotype) is also closely related to Mamu-B*008:01 with a conservative substitution (D101N) at a key F pocket residue in the alpha 1 domain. This asparagine variant in the Mafa-B*008:03 protein is identical to the HLA*B27:03 protein that was also shown to bind a very similar array of peptides as Mamu-B*008:01 (Loffredo et al., 2009). The Mafa-B*008:03 protein also contains a second, nonconservative Y140S substitution at another key F pocket residue in the alpha 2 domain relative to Mamu-B*008:01; this residue is an aspartic acid in HLA-B*27:03 (Loffredo et al., 2009). Fortuitously, the second major class I protein (Mafa-A1*007:05) of the most common MHC-MAFA-NIAID1-000005 haplotype also has a rhesus homologue (Mamu-A1*007:01) with a peptide binding motif that has been determined experimentally (Reed et al., 2011). In this case the Mafa and Mamu protein variants differ at three residues in the alpha 1 and alpha 2 domains, but two of three substitutions are highly conservative. Given the paucity of Mafa class I proteins with known peptide binding motifs, animals that carry the two most common MHC haplotypes in this ICM colony offer a rare resource for studies that could benefit from the ability to monitor CD8 T cell responses. For example, it may be possible to use existing peptide binding motif information to predict epitopes in a virus such as Ebola and then monitor CD8 T cell responses after vaccination and challenge (Sullivan et al., 2011). Moreover, since Mamu-B*008:01 expression is strongly associated with spontaneous control of simian immunodeficiency virus (Loffredo et al., 2009), the high frequency of Mafa-B*008:01 in these ICM could argue against their use in studies where SIV control is an important endpoint.

### Comparison with MHC haplotypes in other cynomolgus macaque populations

As described above, the extended MHC haplotypes that we have characterized in this ICM breeding colony exhibit exceptional diversity. Only two of the 100 extended haplotypes listed in Table 2 appear to have been described previously. Each of the Mafa class I and class II transcripts that are associated with the MHC-MAFA-NIAID1-000056 haplotype are identical to those found on the M3 MHC haplotype of MCM (Budde et al., 2010; O’Connor et al., 2007). Likewise, the MHC-MAFA-NIAID1-000115 haplotype includes class I and class II transcripts that are all identical to the MCM M5 MHC haplotype. Comparison of the STR allele sizes for these two haplotypes with those of MCMs carrying M3 or M5 haplotypes in a separate Alpha Genesis breeding colony revealed identical STR patterns (data not shown). These observations strongly suggest that these are Mauritian-origin MHC haplotypes that were inadvertently introduced to this breeding colony by founders with hybrid ancestry. A review of the pedigree records traced this pair of MCM MHC haplotypes back to two dams (CH2J and CV97) who were included among the original 2002 founders; the sires for both of these breeders are unknown. Similar observations were described for the cynomolgus macaque breeding colony at the BPRC whose founders were reported to have originated in the Indonesian islands and continental Malaysia (Otting et al., 2012). Three of the 32 extended MHC haplotypes characterized in this colony also included class I and class II transcripts that were identical to those that are associated with the M1, M3 or M4 MHC haplotypes of MCMs (Otting et al., 2012; Budde et al., 2010; O’Connor et al., 2007).

The MHC-MAFA-NIAID1-000131 haplotype described here (Table 2) represents a more distant ancestral relationship to the well-characterized M1 haplotype that accounts for nearly 20% of all MHC haplotypes in the MCM population (Wiseman et al., 2013). Although nearly all of the class I transcript sequences associated with this ICM haplotype were identical with the MCM M1 haplotype, several subtle allelic variants were noted. The MHC-MAFA-NIAID1-000131 haplotype is characterized by a major *Mafa-A1*063:03:02* transcript that differs from the *Mafa-A1*063:01* MCM allele by three single nucleotide variants and a single G236E amino acid substitution. Likewise, a single synonymous variant distinguishes the *Mafa-B*104:01:02* and *Mafa-B*104:01:01* transcripts that are associated with these haplotypes. In contrast to the identical STR patterns noted between the MHC-MAFA-NIAID1-000056 and MHC-MAFA-NIAID1-000115 haplotypes versus those of Mauritian-origin individuals, STR allele sizes for the MHC-MAFA-NIAID1-000131 haplotype differ from the M1 MCM haplotype at five of seven STR loci spanning the MHC class I region. In addition, the Mafa class II transcripts for each of these extended MHC haplotypes are completely unrelated. Characterization of MHC class I haplotypes of Filipino cynomolgus macaques by Shiina and coworkers (Shiina et al., 2015) revealed an even more distinct relative of MHC-MAFA-NIAID1-000131 haplotype. The B-Hp2 haplotype which represents one of the most common *Mafa-B* allele combinations observed in of Filipino cynomolgus macaques, also shares allelic variants of five *Mafa-B* lineages (*Mafa-B*104:03, Mafa-B*144:03N, Mafa-B*057:04, Mafa-B*060:02* and *Mafa-046:01:02*) with the Mafa-B104.01.02 regional haplotype that comprises part of the extended MHC-MAFA-NIAID1-000131 haplotype (**Supplemental Figure 5)** as well as the M1 MCM haplotype (Budde et al., 2010). This Filipino B-Hp2 haplotype is also distinguished, however, by at least four completely unrelated *Mafa-B* transcripts (*Mafa-B*050:08, Mafa-B*072:01, Mafa-B*114:02* and *Mafa-I*01:12:01*) relative to its Indonesian and Mauritian counterparts (Shiina et al., 2015).

Broadly speaking, MHC haplotypes that include members of the *Mafa-B*028* and *Mafa-B*021* allele lineages appear to be among the most broadly represented in cynomolgus macaques from different geographic origins as well as related macaque species. In this ICM colony, we identified six distinct variants of Mafa-B028 haplotypes that were associated with six different extended MHC haplotypes (Table 2**, Supplemental Figure 5**). The Mafa-B028.06.01 haplotype described here shares four identical transcripts (*Mafa-B*028:03*, *Mafa-B*021:01, Mafa-B*068:02* and *Mafa-I*01:19*) with Haplotype 23 in the BPRC cynomolgus macaque breeding colony (Otting et al., 2012). While five of six Mafa-B028 haplotypes in the Alpha Genesis colony include a member of the *Mafa-B*124* lineage, the BPRC haplotype was reported to include a *Mafa-B*144:04* transcript. In an earlier study of cynomolgus macaques from several Chinese breeding facilities (Karl et al., 2017), eight distinct Mafa-B028 haplotypes were characterized with different allelic variants of the *Mafa-B*028* and/or *Mafa-B*021* lineages as well as various combinations of additional *Mafa-B* transcripts. Likewise, the B/I-Hp15 haplotype of Filipino cynomolgus macaques was reported to contain yet another combination of *Mafa-B* allelic variants relative to these other breeding centers (Shiina et al., 2015). A variety of Mamu-B028 haplotypes have also been reported in various cohorts of Indian- and Chinese-origin rhesus macaques (Karl et al., 2013; Doxiadis et al., 2013). In addition, PacBio sequencing analyses identified at least three distinct Mane-B028 haplotypes in several pig-tailed macaque cohorts (Semler et al., 2018). Taken together, these observations suggest that the primordial combination of *B*028* and *B*021* genes was likely to have arisen in a common ancestor who predated the divergence of cynomolgus, rhesus and pig-tailed macaques (Smith et al., 2007).

### Concluding remarks

Here, we describe a comprehensive, high-resolution characterization of the *Mafa* class I and class II transcripts that are associated with 100 extended MHC haplotypes in a large, pedigreed colony of ICM. PacBio sequencing of this cohort yielded 202 new *Mafa* transcript sequences with full-length coding regions, bringing the number in IPD-MHC Release 3.3.0.0 (2019-06-13) build 126 to a total of 2,489 *Mafa* sequences. This study was informed by an extensive database of STR patterns for a panel of markers that span the 5 Mb MHC genomic region that have been collected since this breeding colony was established in 2002. The STR data allowed us to define the allele content of extended MHC haplotypes for individuals that were known to share chromosomes that were identical by descent rather than simply inferring haplotypes based on combinations of class I and class II transcripts that appear to travel together in multiple animals who lack known pedigree relationships. A large majority of the 70 Mafa-A, 78 Mafa-B and 45 Mafa-DRB haplotypes identified here were only associated with one or two of the 100 extended MHC haplotypes that we characterized. This extensive MHC diversity was derived from only approximately 79 founding animals for this breeding colony. In contrast, a comparable study of Indian-origin rhesus only identified 17 Mamu-A, 18 Mamu-B and 22 Mamu-DRB haplotypes in the BPRC breeding colony that was established with 137 founding animals (Doxiadis et al., 2013).

Although this dataset is restricted to a single ICM breeding colony, it provides a glimpse of extraordinary MHC diversity that must be present in the wild population of ICM as a whole. It is reasonable to predict that the high level of genetic diversity described here for the MHC region will extend across the rest of the ICM genome. As previously noted, MCM are thought to be derived from a founder population of ICM (Lawler et al., 1995; Tosi and Coke, 2007) which significantly limited the genomic diversity in MCM, especially in the MHC region where only seven ancestral haplotypes are observed. While this restricted genetic diversity is valuable in many areas of research, it can present a challenge for transplant investigators whose goal is to maximize MHC disparity between donor and recipient tissues. The ICM population, in contrast, exhibits considerably more diversity, increasing its value as a model species for biomedical research, such as transplantation studies. This pedigreed cynomolgus macaque cohort also provides a valuable resource for future studies designed to characterize additional immune-important genomic regions such as the killer immunoglobulin-like receptors (Bimber and Evans, 2015; Prall et al., 2017; Bruijnesteijn et al., 2018) and Fc gamma receptors (Haj et al., 2019).

---

**Supplemental Figure 1:** Selected demographic information for Alpha Genesis Indonesian cynomolgus macaques investigated in this study. Animal IDs are shown in orange for female macaques and blue for male macaques.

**Supplemental Figure 2:** STR patterns for extended MHC haplotypes. The values listed here are the length of the PCR products for each of the eleven STR loci spanning the MHC region that were associated with each haplotype.

**Supplemental Figure 3:** Primer sequences used for PCR amplification of cDNA templates for MHC class I and II.

**Supplemental Figure 4:** Regional haplotype definitions for Mafa-A. In the leftmost column, green indicates direct observation of inheritance; gray indicates that no direct inheritance was observed. Gray italics indicate an inferred allele.

**Supplemental Figure 5:** Regional haplotype definitions for Mafa-B. In the leftmost column, green indicates direct observation of inheritance; gray indicates that no direct inheritance was observed. Gray italics indicate an inferred allele.

**Supplemental Figure 6:** Regional haplotype definitions for Mafa-DRB.

**Supplemental Figure 7:** PacBio sequencing results for MHC class I and II for 295 ICM.

**Supplemental Figure 8:** MiSeq sequencing results for MHC class I and II for 295 ICM.

## Acknowledgements

We appreciate pedigree information provided by Isabelle Lussier for the Alpha Genesis ICM breeding colony. We also thank the University of Wisconsin Biotechnology Center DNA Sequencing Facility for providing Pacific Biosciences library preparation and sequencing services. In addition, we are extremely grateful for the expert assistance of Nel Otting and Natasja de Groot in providing official IPD-MHC allele nomenclature for the novel sequences and extensions of partial sequences that are reported here. This work was made possible by financial support through contract HHSN272201600007C from the National Institute of Allergy and Infectious Diseases of the NIH. This work was also supported in part by the Office of Research Infrastructure Programs/ OD (P51OD011106) awarded to the Wisconsin National Primate Research Center at the University of Wisconsin-Madison. This research was conducted in part at a facility constructed with support from Research Facilities Improvement Program grants RR15459-01 and RR020141-01.

